# Whole-brain analysis of CO_2_ chemosensitive regions and identification of the retrotrapezoid and medullary raphé nuclei in the common marmoset (*Callithrix jacchus*)

**DOI:** 10.1101/2023.09.26.558361

**Authors:** Ariana Z. Turk, Marissa Millwater, Shahriar SheikhBahaei

**Affiliations:** Neuron-Glia Signaling and Circuits Unit, National Institute of Neurological Disorders and Stroke (NINDS), National Institutes of Health (NIH), Bethesda, 20892 MD, USA

**Keywords:** Astrocyte, CO_2_ chemosensitivity, common marmoset, glia, retrotrapezoid nucleus, medullary raphé

## Abstract

Respiratory chemosensitivity is an important mechanism by which the brain senses changes in blood partial pressure of CO_2_ (*P*CO_2_). It is proposed that special neurons (and astrocytes) in various brainstem regions play key roles as CO_2_ central respiratory chemosensors in rodents. Although common marmosets (*Callithrix jacchus*), New-World non-human primates, show similar respiratory responses to elevated inspired CO_2_ as rodents, the chemosensitive regions in marmoset brain have not been defined yet. Here, we used c-fos immunostainings to identify brain-wide CO_2_-activated brain regions in common marmosets. In addition, we mapped the location of the retrotrapezoid nucleus (RTN) and raphé nuclei in the marmoset brainstem based on colocalization of CO_2_-induced c-fos immunoreactivity with Phox2b, and TPH immunostaining, respectively. Our data also indicated that, similar to rodents, marmoset RTN astrocytes express Phox2b and have complex processes that create a meshwork structure at the ventral surface of medulla. Our data highlight some cellular and structural regional similarities in brainstem of the common marmosets and rodents.

## Introduction

Carbon dioxide (CO_2_) respiratory chemosensitivity is the mechanism to adjust breathing in response to changes in brain parenchymal partial pressure of CO_2_ (*P*CO_2_) and is a key regulatory mechanism of breathing in mammals. Specialized cells located in the carotid bodies and brainstem are responsible for this respiratory response (Heymans and Bouckaert 1930; Heymans and Neil 1958). The current model of central respiratory chemosensitivity hypothesizes that groups of specialized brainstem neurons in the retrotrapezoid nucleus (RTN) and medullary raphé nucleus are responsible for central pH/CO_2_ sensing and adjusting ventilation (Teran et al. 2014; Guyenet et al. 2019), though, other

Chemosensitive region of RTN is located ventral to the facial nucleus in the medulla oblongata (Smith et al. 1989; Del Negro et al. 2018; Nattie and Li 1994). Increases in CO_2_/H^+^ robustly activate chemosensitive neurons and astrocytes in the RTN (Mulkey et al. 2004; Wang et al. 2013; Onimaru et al. 2012; Sobrinho et al. 2017; Czeisler et al. 2019; Erlichman and Leiter 2010; Gourine et al. 2010) and induces expression of immediate early gene (IEG, such as c-fos) in RTN cells in rodents and cats (Teppema et al. 1997; Sato et al. 1992; Teppema et al. 1994). In rodents, RTN neurons and astrocytes have been shown to respond to hypercapnia (increased inspired CO_2_) and acidosis, and adjust the respiratory drive (Wang et al. 2013; Kumar et al. 2015; Guyenet et al. 2019; Guyenet et al. 2009; Stornetta et al. 2006; Gourine et al. 2010; Korsak et al. 2018; Marina et al. 2018). In addition, it is proposed that the RTN integrates chemosensory information and may act as a critical hub for respiratory chemoreception (Guyenet et al. 2019; Ruffault et al. 2015). In rodents, neurons (and some astrocytes) in this region are derived from dorsal progenitor cells that express paired-like homeobox 2B (*Phox2b*), (Czeisler et al. 2019; Dubreuil et al. 2009). Accordingly, Phox2b has been used as a marker of RTN neurons in adult rats to investigate central chemoreflexes (Abbott et al. 2011; Kang et al. 2007; Takakura et al. 2014; Abbott et al. 2013; Stornetta et al. 2006; Marina et al. 2010; Wang et al. 2013; Ruffault et al. 2015; Ramanantsoa et al. 2011). In humans, mutations of *PHOX2B* cause central congenital hypoventilation syndrome (CCHS) (Dubreuil et al. 2008; Weese-Mayer et al. 2005), which is characterized by impaired breathing control and failures in chemosensitivity.

In rodent models, it is also suggested that activity of serotonergic (5-HT) neurons is critical to central chemosensitivity (Iceman et al. 2013; Corcoran et al. 2013). Rodents display perturbed ventilation, especially impaired ventilatory response to hypercapnia, when 5-HT function is altered in neurons and astrocytes in the raphé nuclei by genetic tools, pharmacological lesioning, or selective silencing (Ikoma et al. 2018; Wang et al. 2001; Wang et al. 1998; Wu et al. 2019; Patodia et al. 2019; Duncan et al. 2010; Massey et al. 2015; Nattie 1999; Kiyasova et al. 2011; Iceman et al. 2013; Buchanan and Richerson 2010; Hilaire et al. 2010; Hodges et al. 2008; Hodges et al. 2009; Hodges and Richerson 2010; Nattie and Li 2009; Nattie et al. 2004; Ray et al. 2011). In these serotonergic neurons, tryptophan hydroxylase (TPH) converts L-tryptophan to 5-HT. Although many cells in the brain can accumulate and release 5-HT, TPH expression is specific for the 5-HT-producing neurons, and therefore TPH is also used as a marker to identify 5-HT neurons (Gaspar and Lillesaar 2012; Gaspar et al. 2003). In mammals, central 5-HT is confined within the raphe nuclei brainstem, and hypercapnia induces 5-HT release in respiratory motor nuclei (Harper et al. 2005) as well as expression of IEG in raphé serotonergic neurons (Pete et al. 2002; Johnson et al. 2005; Larnicol et al. 1994).

In addition to neurons, brainstem astrocytes in ventrolateral medulla have been proposed to have a key role in central CO_2_ chemosensitivity (Turovsky et al. 2016; Beltrán-Castillo et al. 2017; Eugenín León et al. 2016; Gourine et al. 2010; Erlichman and Leiter 2010; Mulkey and Wenker 2011; Huckstepp et al. 2010; Sheikhbahaei, Turovsky, et al. 2018; van de Wiel et al. 2020). For instance, RTN astrocytes create a meshwork of dense processes at the RTN (Sheikhbahaei, Morris, et al. 2018) and respond to changes of pH by releasing adenosine triphosphate (ATP) (Erlichman and Leiter 2010; Gourine et al. 2010; Huckstepp et al. 2010).

Recently we have shown that hypercapnia augments frequency and depth of breathing in the common marmoset (*Callithrix jacchus*) (Bishop et al. 2021), a New World non-human primate animal model. This hypercapnia-induced respiratory response is comparable to respiratory responses reported in other mammals (Boddy et al. 1974; Hoffman et al. 1982; Kuwaki et al. 1996; Douglas et al. 1982; Kong and Berger 1986). However, the chemosensitive regions in the marmoset’s brain have not been mapped. Accordingly, in this study, we aimed to map central chemosentive regions of the RTN and medullary 4aphe in the common marmoset brainstem by describing the expression of IEG activation as well as Phox2b and TPH immunostainings. The description of Phox2b and TPH expression within the brainstem of the marmoset is likely to help to locate the central chemosensetive regions. In addition, measuring IEG activations, we also identified other brain regions, in which neuronal activities may be increased by systemic hypercapnia.

## 2. Materials and Methods

### 2.1 Animals

Four adult common marmosets (*Callithrix Jacchus*) (2 males and 2 females; average age of 60 ± 6.5 months; average weight of 402 ± 19.5 g) were used in this study. Marmosets were housed in cages in pairs or alone in a room with a 12h light/dark cycle. Their food and water intake were regulated, receiving food and water ad liberum. We complied with all relevant ethical regulations for animal testing and research. All procedures in this study were approved by the Animal Care and Use Committee of the Intramural Research Program of National Institute of Mental Health (NIMH).

### 2.2 Exposure of marmosets to hypercapnia by whole body plethysmography

Whole-body plethysmography was used to record respiratory activity in unrestrained awake marmosets, as described previously (Bishop et al. 2021). Briefly, on the day of the experiment, the marmoset was placed in a Plexiglas recording chamber (∼3 L) that was flushed continuously with a humidified room air (temperature 26–28 °C) at a rate of 2.2 L min^−1^. The experimental animals (1 male and 1 female) were allowed to acclimate to the chamber environment for at least 30 min before application of 10 min hyperoxic hypercapnia (60% O2, 6% CO2, balanced with nitrogen). Animals were sacrificed 1.5 hrs after the hypercapnic experiment. The control animals (1 male and 1 female) were not exposed to hypercapnia.

### 2.3 Tissue Processing and Immunohistochemistry

Marmosets were euthanized with an overdose of anesthesia sodium pentobarbital (100 mg kg^−1^, i.p.) and transcardially perfused with 500 mL of phosphate-buffered saline (PBS, 0.1M) solution followed by 4% paraformaldehyde (PFA) fixative. Subsequently, the brain was extracted and post-fixed for 3 to 5 days in the same PFA solution.

Ex-vivo MRI were performed on the extracted brains, in which the brains were scanned on a 7T/30-cm horizontal bore MRI spectrometer (Bruker Biospin) with a 30-mm inner diameter quadrature millipede coil (ExtendMR). The brains were then sectioned serially at 50 µm. Floating slices were divided into ten series and stored in anti-freeze solution in −20 °C until staining. Coronal sections were immunostained as described before (Sheikhbahaei, Morris, et al. 2018; Turk and SheikhBahaei 2021). Briefly, the brain sections (1 every 10 sections) were incubated on a shaker for 48 h at 4 °C with a primary antibody. While some primary antibodies were conjugated and did not require a secondary antibody, other sections were incubated with primary antibodies that required secondary conjugation (see Table 1 for more information). Those sections were subsequently incubated in secondary antibodies conjugated to the fluorescent probes (1:500; Life Technologies) for 1.5 h shaking at room temperature. A separate section before or after the one that used for antibody staining was stained with Nissl, as described before (Turk and SheikhBahaei 2021). Briefly, the tissue was stained with NeuroTrace™ green fluorescent Nissl stain (1:200 in PBS, Life Technologies, catalog No. N-21480) for 20 min at room temperature. These Nissl-stained sections and MRI images were used to put the sections in order and mapped to the corresponding sections in marmoset atlas (Paxinos et al. 2012). All sections were subsequently mounted onto microscope slides and covered in an anti-fading mounting media. Tiled images were acquired using an AxioScan.Z1 slide scanner (Carl Zeiss, with a 20X objective) and used for quantification of immunostained cells. Using an inverted confocal laser scanning microscope (Zeiss LSM 510) with image acquisition settings with 1024 x 1024-pixel resolution, z-stack images of Phox2B, TPH, GFAP, and c-fos-positive cells were obtained from the slice thickness at 20x and 40x in various brain regions. To avoid possible experimental variation, immunostaining was conducted and processed by one investigator with the same solutions and imaging protocol.

**Table 1.** Primary Antibody Characterization.

| Antigen | Description of Immunogen | Source, host species, catalog No. RRID | Dilution used |
| --- | --- | --- | --- |
| c-fos | Synthetic peptide corresponding to amino acids 2 through 17 from rat c-fos. Subtype IgG2a ($\kappa$ light chain) | Synaptic Systems, rat monoclonal, catalog #226017, RRID: AB_2864765 | 1:250 |
| c-fos | Synthetic peptide corresponding to amino acids 2 through 17 from rat c-fos | Synaptic Systems, guinea pig polyclonal, catalog #226004, RRID: AB_2619946 | 1:250 |
| Phox2b | 14 amino acid C-terminal sequence of Phox2B protein | Gift from J-F. Brunet, Institut de Biologie de l'École Normale Supérieure, Paris, FR; RRID: AB2315161 | 1:500 |
| TPH | Purified protein A/G | Sigma-Aldrich, mouse monoclonal, catalog # AMAAB91108, RRID: AB_2665804 | 1:200 |
| GFAP | GFAP isolated from cow spinal cord | DAKO, rabbit polyclonal, catalog #z-0334, RRID: AB_10013382 | 1:1,000 |
| GFAP | Purified glial filament protein | Sigma-Aldrich, mouse monoclonal Cy3 conjugate, MAB3402C3, RRID: AB_11213580 | 1:1,000 |

### 2.4 Antibody Characterization

We used anti-Phox2B and anti-TPH antibodies to identify RTN and raphé nucleus, respectively. Anti-Phox2B polyclonal rabbit antibody (1:500, gift from J-F. Brunet, Institut de Biologie de l’École Normale Supérieure, Paris, FR; RRID: AB2315161) was raised against the 14-AA C-terminal sequence of the Phox2B protein. The rat and mouse sequence are identical, with the one addition of tyrosine at the N-terminal in the rat sequence. Specificity of this antibody was confirmed via in situ hybridization and in a Phox2B knock-out mouse model (Pattyn et al. 1997). This antibody has also previously been used in a non-human primate model (Levy et al. 2019).

An anti-tryptophan hydroxylase 2 (anti-TPH2) mouse monoclonal antibody 1:200, Sigma-Aldrich, catalog #AMAAB91108, RRID: AB_2665804) was used for dorsal raphé identification. Tryptophan hydroxylase is a serotonin (5-HT) precursor, particularly enriched in neurons of the pons and medulla in human (Zhang et al. 2004). This specific antibody was developed and validated by the Human Protein Atlas (HPA) project (www.proteinatlas.org) where it was shown to be enriched in neuronal cell bodies, projection and synapses. Cross-reactivity has been shown with human, rat, and mouse tissue and has reviously been used in rodent and human tissue (Lopez et al. 2014; Lentini et al. 2021).

We used histology techniques to count the number of cells with activated Immediate Early Genes (IEG; c-fos) to identify the regions activated by hypercapnic challenge (including CO_2_ chemosensitive regions). IEG activate rapidly and transiently by various extracellular stimuli, including CO_2_ (Okada et al. 2002; Wickström et al. 2002; Haxhiu et al. 1996). Here, we used c-fos to identify brain cells and regions that were activated by hypercapnic effects. Two anti-c-fos antibodies were used to validate specificity of c-fos staining in the adult marmoset (see Table 1). These antibodies have previously been successfully used as a marker for neuronal activity in rodent, non-human primate, and human tissue (Sarkar et al. 2021; Heukelum et al. 2021; Lopez et al. 2014). The polyclonal guinea pig anti-cFps antibody (1:250, Synaptic Systems, catalog #226004, RRID: AB_2619946) has a synthetic peptide corresponding to amino acids 2 to 17 of rat c-fos. This antibody has been shown to show reactivity in human, mouse, rat, monkey, ape, dog, and bovine (manufacturer’s technical information). The second c-fos antibody used was a monoclonal rat anti-c-fos antibody (1:250, Synaptic Systems, catalog #226017, RRID: AB_2864765). Similar to the above-mentioned c-fos antibody, this too holds a synthetic peptide corresponding to amino acid 2 to 17 of rat c-fos with a subtype 1gG2 (k light chain). This antibody reacted specifically with c-fos in immunoblotting assays with a single band found at 60kDa (manufacturer’s technical information).

To identify astrocytes, we used antibodies against glial fibrillary acidic protein (GFAP). GFAP is isolated from cow spinal cord, the rabbit-polyclonal anti-GFAP antibody (1:1,000; Agilent (formerly DAKO), catalog #z-0334, RRID: AB_10013382) cross-reacts with an epitome of mouse, rat, and human cytoskeleton, the intra-cytoplasmic filamentous protein [manufacturer’s technical information; (Eng 1985; Eng et al. 2000; Turk and SheikhBahaei 2021)]. Additionally, this antibody stains a double band at 245-395kDA on Western Blot Analysis (Key et al. 1993). We have recently showed that this antibody also reacts with marmoset’s astrocytes (Turk and SheikhBahaei 2021).

The mouse monoclonal GFAP antibody (1:1,000, Sigma-Aldrich, catalog #MAB3402C3, RRID: AB_11213580) has the ability to react with human, pig, chicken, and rat GFAP (manufacturer’s technical information). This GFAP antibody was raised against purified glial filament protein (Debus, Weber, & Osborn, 1983). We have previously shown the specificity of this antibody to detect astrocytes in marmoset tissue as well (Turk and SheikhBahaei 2021).

### 2.5 Cell count

Pipsqueak AI^TM^ (Rewire Neuro, Inc., RRID:SCR_022149) was used to unbiasedly obtain a cell count of neurons from each region of interest (Eisele et al. 2021). Images were imported into ImageJ where the Pipsqueak AI extension was then used. Single label analysis of neurons from each region were obtained. To normalize and properly compare the regions, the same volumetric size was used for each region expressed at density/mm^2^. C-fos count was normalized to the total number of c-fos positive cells in each animal (Kakall et al. 2019; Biro et al. 2017).

### 2.6 Statistical analyses

The data were analyzed in Prism 9.0 software (Graphpad Software Inc., RRID: SCR_002798) and reported as averages ± standard error of mean (SEM). Unpaired *t* tests were used for statistical analyses between control and experimental groups.

## 3. Results

### 1. Brain-wide responses to acute hypercapnia

We first performed a whole-brain survey of c-fos expression in cortical, midbrain and brainstem regions in marmosets exposed to hypercapnia. We compared c-fos expression in the brainstem in animals exposed to either room air (0% CO_2_; control group; *n* = 2) or acute hypercapnia (6% CO_2_; experimental group; *n* = 2) and found no difference between the two groups in a hypercapnic state compared to control (p = 0.1, 0.5 ± 0.02 vs. 0.4 ± 0.01 in control, unpaired *t* test) (Figure 1, Table 2). In the midbrain, we observed an increase in c-fos expression in the ventrolateral periaqueductal grey (VLPAG) (p = 0.02, 0.07 ± 0.009 vs. 0.01 ± 0.0004 in control, unpaired *t* test) in marmosets exposed to CO_2_. In the hypothalamic regions, exposure to CO_2_ only induced higher c-fos expression in the lateral hypothalamus (p = 0.03, 0.2 ± 0.03 vs. 0.02 ± 0.004 in control, unpaired *t* test), but not the posterior or ventromedial hypothalamic areas (p = 0.5, 0.09 ± 0.01 vs. 0.08 ± 0.003 in control; p = 0.09, 0.06 ± 0.01 vs. 0.02 ± 0.0003 in control, respectively, unpaired *t* test). Increased in c-fos expressions were also observed in the paraventricular nucleus of the thalamus (PVT) in marmosets exposed to CO_2_ condition (p = 0.067, 0.05 ± 0.01 vs. 0.03 ± 0.0003 in control). In evaluating the basal ganglia region, we found no differences in c-fos expression in ventral tegmental area (VTA) (p = 0.4, 0.03 ± 0.009 vs. 0.02 ± 0.002 in control, unpaired *t* test) and substantia nigra (SN) (p = 0.4, 0.03 ± 0.01 vs. 0.02 ± 0.009 in control, unpaired t test) between the marmosets exposed to 6% CO_2_ or 0% CO_2_. However, hypercapnia increased c-fos expression in the caudate (p = 0.07, 0.1 ± 0.006 vs. 0.07 ± 0.01 in control, unpaired *t* test). We also looked at the c-fos expression in central nucleus of the amygdala (CeA) and found no expression differences between hypercapnic and control groups (p = 0.1, 0.07 ± 0.004 vs. 0.06 ± 0.002 in control, unpaired *t* test). In cortical regions, there were differences in c-fos expression in the insula (p = 0.006, 0.07 ± 0.03 vs. 0.005 ± 0.003 in control, unpaired *t* test) and in area 24 (p = 0.002, 0.07 ± 0.002 vs. 0.02 ± 0.0004 in control, unpaired *t* test) while no differences were seen in the other cortical regions (Figure 1).

**Figure 1:**
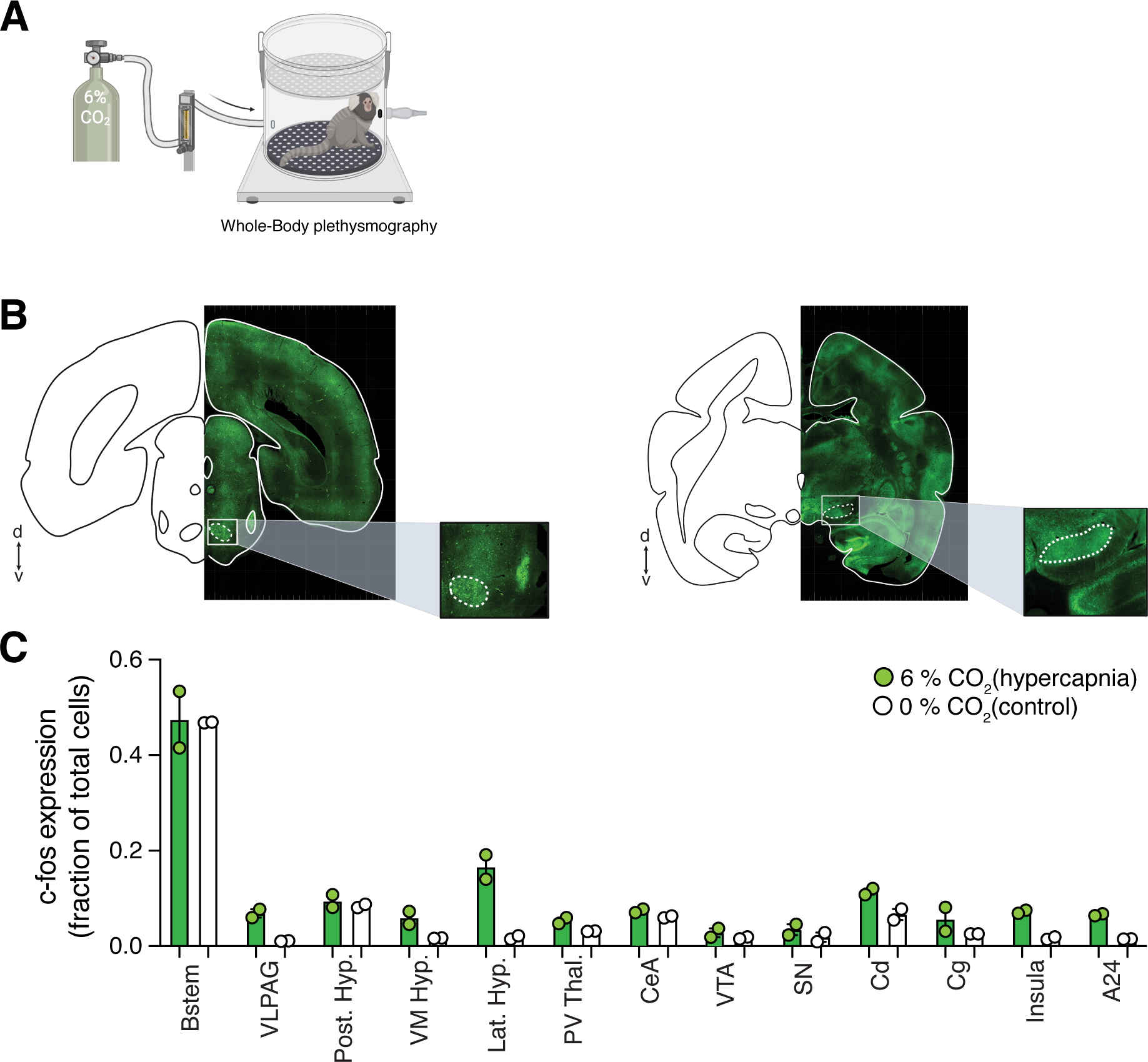
Whole-brain analysis of c-fos expression following hypercapnic exposure. Illustrated image of marmoset whole-body plethysmography experiment with 6% CO_2_ **(a)**. Traces and whole brain images with examples of c-fos positive regions enlarged **(b)**. C-fos expression in regions of interest examined in animals exposed to 6% CO_2_ and normal CO_2_ levels **(c)**. Brainstem (Bstem), ventrolateral periaqueductal gray (VLPAG), posterior hypothalamus (post. Hyp.), ventromedial hypothalamus (VM hyp.), paraventricular thalamus (PV Thal.), ventral tegmental area (VTA), central nucleus of the amygdala (CeA), substantia nigra (SN), caudate (Cd), cingulate (Cg), Insula, area 24 (A24).

**Table 2.** Brain-wide c-fos expression in marmosets exposed to acute hypercapnia (6% CO_2_)

|  | **6% CO<sub>2</sub>**<br>**(mean±SEM, n = 2)** | **0% CO<sub>2</sub>**<br>**(mean±SEM, n = 2)** | ***p* value**<br>**(unpaired *t***<br>**test)** |
| --- | --- | --- | --- |
| Brainstem | 0.5±0.02 | 0.4±0.01 | 0.1 |
| Ventrolateral<br>Periaqueductal Grey<br>(VLPAG) | 0.07±0.01 | 0.011±0.0004 | 0.022 |
| Posterior<br>Hypothalamus | 0.09±0.01 | 0.08±0.003 | 0.5 |
| Ventromedial<br>Hypothalamus | 0.06±0.01 | 0.02±0.0003 | 0.09 |
| Lateral Hypothalamus | 0.16±0.03 | 0.02±0.004 | 0.03 |
| Paraventricular<br>nucleus of the<br>thalamus (PVT) | 0.05±0.01 | 0.03±0.0003 | 0.067 |
| Ventral Tegmental<br>Area (VTA) | 0.03±0.01 | 0.02±0.002 | 0.4 |
| Central nucleus of the<br>Amygdala (CeA) | 0.07±0.003 | 0.06±0.002 | 0.1 |
| Substantia Nigra (SN) | 0.03±0.01 | 0.02±0.01 | 0.4 |
| Caudate nucleus | 0.11±0.01 | 0.07±0.01 | 0.076 |
| Posterior cingulate<br>cortex | 0.06±0.02 | 0.03±0.0001 | 0.3 |
| Anterior cingulate<br>cortex (ACC) | 0.07±0.002 | 0.02±0.0004 | 0.002 |
| Insula cortex | 0.07±0.003 | 0.01±0.003 | 0.006 |

### 2. Putative RTN region in marmoset brainstem

In marmoset brain, most of the CO_2_-induced c-fos-positive cells were in the brainstem (Figure 2). In rodents and humans, RTN and raphé nucleus are proposed to act as central CO_2_ chemosensory regions (Guyenet et al. 2009; Teran et al. 2014). Therefore, we mapped these regions in marmoset brainstem using histology techniques.

**Figure 2:**
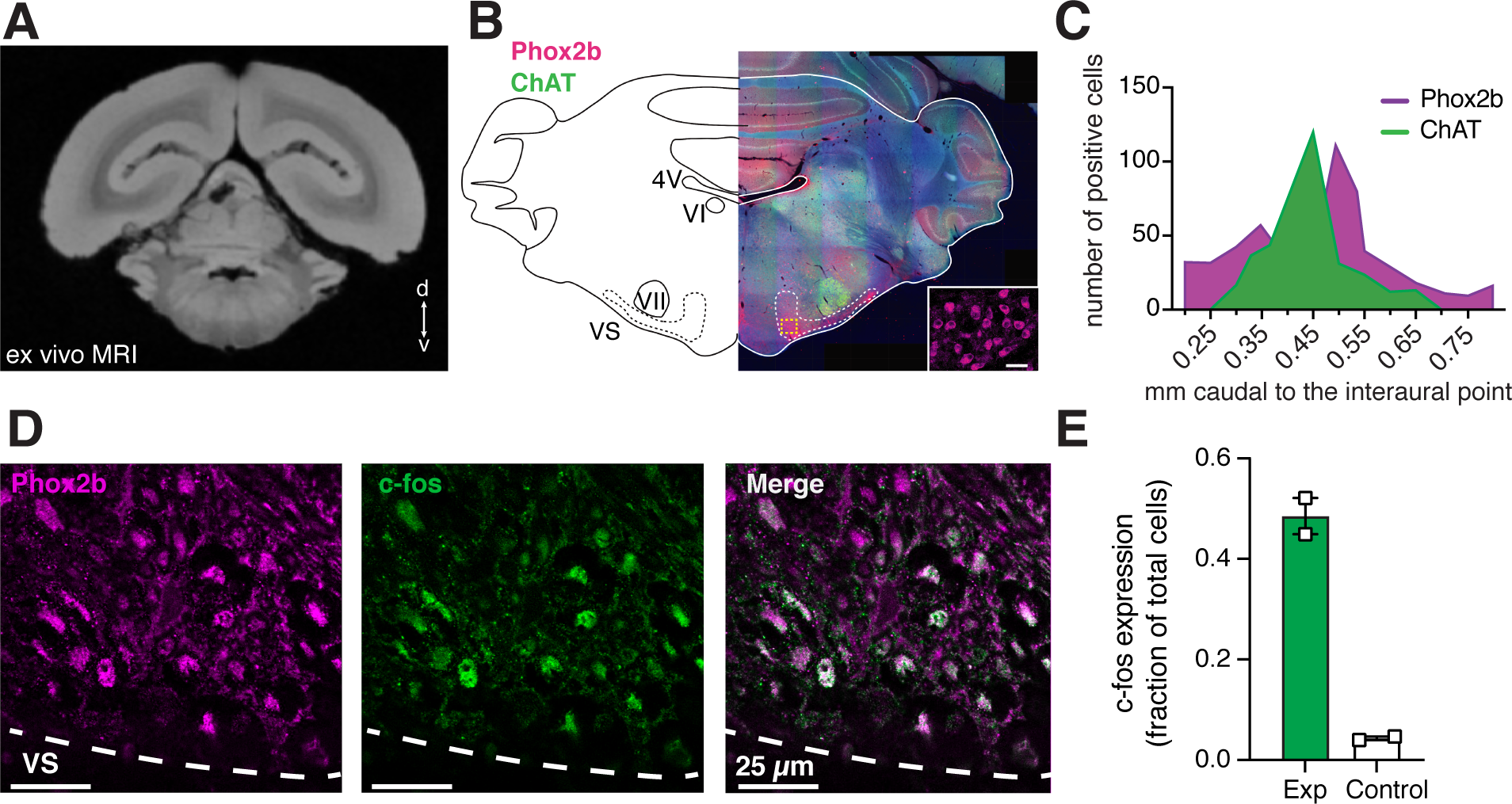
Identification of retrotrapezoid nucleus. Ex vivo coronal MRI image of marmoset brainstem slice including the retotrapezoid nucelus (RTN) **(a)**. Whole-slice brain image of brainstem plane that includes the RTN and overlying trace. Bottom right demonstrates high magnitude image of Phox2b^+^ cells in the RTN **(b)**. A histogram demonstrating the peak location of Phox2b^+^cells which represents the location of the RTN is shown in purple and ChAT-positive cells representing the facial nucleus are illustrated in green **(c)**. Phox2b^+^ cells in purple are colocalized with cfos-positive cells in green within the RTN **(d)**. Cfos expression was quantified in the RTN on animals under 6% CO2 conditions and normal conditions (p = 0.006, Mann-Whitney t-test)(e).

Phox2b-positive cells of RTN are located ventral to the facial motor nucleus in rodents and primates (Kang et al. 2007; Abbott et al. 2011; Ruffault et al. 2015). Therefore, we immunostained serial brainstem sections with antibodies against Phox2b and Choline acetyltransferase (ChAT, which was used as a marker for facial motor nucleus, Table 1) (Figure 2A and 2B). We used MRI data to align sections in order and identified the peak facial nucleus location at – 0.45 mm from the interaural point (Figure 2C). Location of RTN was depicted in a histogram quantifying the number of ChAT-positive neurons (i.e., facial nucleus) and Phox2b**^+^** cells. To verify the RTN location, we also quantified the c-fos and phox2b expression in the marmosets exposed to hypercapnia (n = 2) and control groups (n = 2) (Figure 2D-F) within the putative RTN region (Figure 2G). When comparing the c-fos expression, marmosets exposed to acute CO_2_ illustrated significantly higher numbers of c-fos expression in Phox2b**^+^** cells in the parafacial regions (p = 0.007, 0.5 ± 0.04 vs. 0.04 ± 0.004 in control, unpaired *t* test; Figure 2G). We estimated that there are 2543 ± 174 Phox2b**^+^** c-fos**^+^** neurons in the marmoset putative RTN region.

### 3. Putative medullary raphé nuclei in marmoset brainstem

We then used similar methods to locate the raphé nuclei in marmoset brainstem. To do so, we used immunostaining against tryptophan hydroxylase (TPH, which is a serotonin precursor used to identify 5-HT-producing neurons within the raphé nucleus, Table 1; Figure 3A). Because raphé nuclei transcends a large portion of the brainstem, we quantified TPH**^+^** cells over larger brainstem sections and identified the raphé nuclei at all levels, including dorsal raphé and raphé obscurus (Figure 3B). We also found high colocalization of TPH neurons and c-fos expression in medullary raphé as well dorsal raphé nucleus (DRN), which further confirms that these putative raphé neurons are in fact activated by systemic CO_2_ (Figure 3C-E).

**Figure 3:**
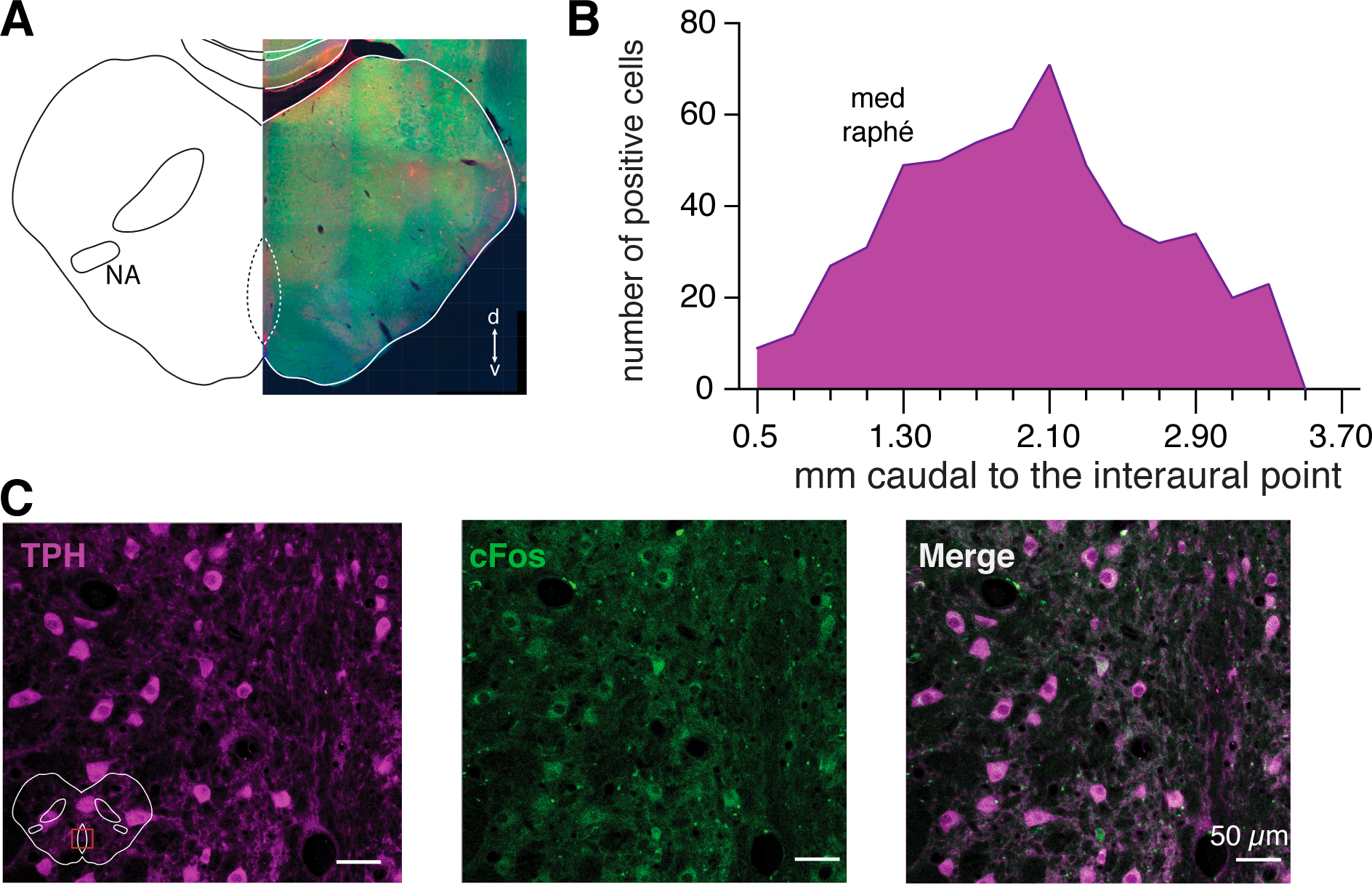
Identification of c-fos-expressing cells in the raphé nucleus. Whole-brain image of a plane including a portion of the raphé nucleus along with a trace **(a)**. The histogram in **(b)** illustrates the increased expression in TPH which peaks in medial raphé nucleus. Immunofluorescence demonstrates colocalization of TPH and c-fos within the raphé nucleus **(c)**.

### 4. Brainstem astrocytes

Astrocytes have also been proposed to act as CO_2_ chemosensors in the ventrolateral medulla, including RTN region (Gourine et al. 2010). Accordingly, we investigated astrocytes morphology in RTN and medullary raphé nucleus. We found dense astrocytic processes within the RTN region both in the ventral (Figure 4A) and lateral regions (Figure 4B). Interestingly, while the processes in the lateral RTN region are perpendicular to the ventral surface, the astrocytic processes in the ventral RTN region are parallel to the ventral surface (Figure 4A, 4B). In addition to RTN region, our data suggest dense astrocytic processes in the raphe nucleus (Figure 4C). In rodents, Phox2b**^+^** astrocytes have been reported in RTN (Czeisler et al. 2019). Consistent with these data, we also found that some RTN astrocytes express Phox2b (Figure 5).

**Figure 4:**
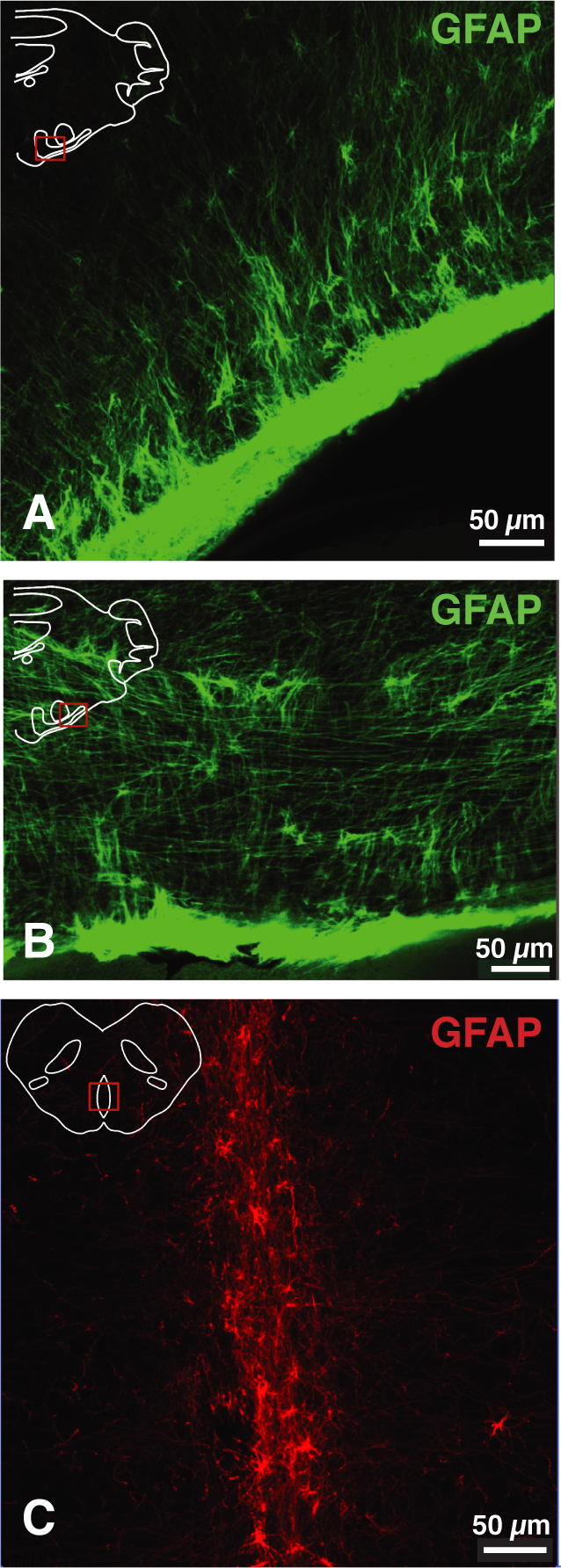
GFAP staining in brainstem regions. GFAP staining on the ventromedial surface of the brainstem exhibit astrocytic processes perpendicular to the ventral surface **(a)** while GFAP staining on the ventrolateral surface in the same plane demonstrated astrocytic fibers parallel to the ventral surface **(b)**. Dense astrocytic processes were also observed in the raphé nucleus **(c)**.

**Figure 5:**
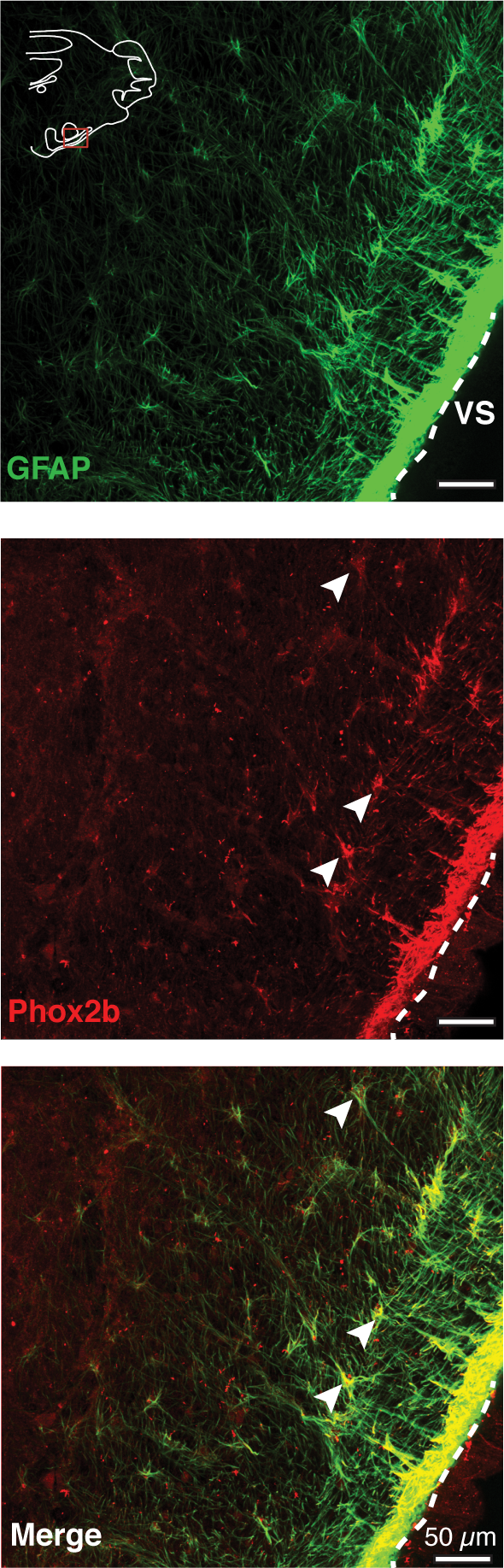
GFAP-Phox2B colocalization in the RTN. GFAP-positive astrocytes (green) and Phox2b^+^ staining (red) on the ventral surface (VS) of a brainstem slice that is identified as the retrotrapezoid nucleus. The colocalization is illustrated in merged image on the bottom.

## 4. Discussion

Breathing behaviors and the mechanisms that regulate respiratory activity are highly preserved among mammalian species (Milsom 2002; Glass and Wood 2009). The neural circuits for control of breathing are compartmentalized within regions in the medulla and pons (Smith et al. 2009; Del Negro et al. 2018). These centers receive direct and indirect modulatory inputs from O_2_ and CO_2_ chemosensors for homeostatic control of blood partial pressure of O_2_ (*P*O_2_) and *P*CO_2_/pH. Data from rodents and cats suggest that RTN and medullary raphé (i.e., raphe magnus and pallidus) act as central CO_2_ chemosensitive center in the brainstem (Stornetta et al. 2006; Teppema et al. 1994; Takakura et al. 2014). While it has been shown that the common marmoset, a New-World non-human primate, has a similar hypercapnic respiratory response to that of rodents and humans (Bishop et al. 2021), the brainstem central chemosensitive regions of the RTN and medullary raphé have yet to be mapped. Accordingly, in this study we performed a whole-brain analysis of CO_2_ chemosensitive regions in marmoset’s brain and additionally identified putative locations of RTN and medullary raphé in the marmoset brainstem.

### 4.1 c-fos immunohistochemistry

Immediate Early Genes (IEGs), such as *c-fos*, is a widely tool that has been used to map rapidly activated neurons by extracellular stimuli (Herdegen and Leah 1998; Bullitt 1990). The expression of the IEGs is considered to be a marker for activation of individual neurons after influx of Ca^2+^ influx through NMDA receptors and L-type voltage-sensitive Ca^2+^ channels (VSCC) (Liste et al. 1995; Berretta et al. 1997; Parthasarathy and Graybiel 1997; Murphy et al. 1991). Rapidly and transiently induced c-fos, the nuclear protein Fos, can be identified by immunofluorescent techniques (Kobelt et al. 2004; Perrin-Terrin et al. 2016; Ceccatelli et al. 1989). Several studies have used antibodies against c-fos to map neuronal cells involved in the physiological response to hypercapnia (Teppema et al. 1997; Pete et al. 2002; Sato et al. 1992; Teppema et al. 1994). This histology method has been used in common marmosets as well to illustrate activation of neuronal cells by various stimuli (Magalhães et al. 2021; Barros et al. 2015; Miller et al. 2010). Compared to rodents, c-fos expression in marmosets is more broadly expressed (5× more cells) and the expression lasts longer (up to 3 hr.) (Magalhães et al. 2021). Accordingly, we employed c-fos immunostaining to identify brain regions that are activated by acute hypercapnia (Figure 1).

We identified a higher number of c-fos**^+^** cells in the brainstem compared to the rest of the brain, even in the control marmosets that were not exposed to hypercapnia. This could be because brainstem neurons are highly active at the baseline to support vital physiological activities, including control of cardio-respiratory system. This basal c-fos activity has been reported in cats and rodents as well (Joubert et al. 2016; Morales et al. 1999; Sato et al. 1992).

### 4.2. CO_2_ chemosensitive brain region

We found higher c-fos expression in certain brain regions in marmosets exposed to acute 6% CO_2_ compared to marmosets that breathed room air. These regions include ventrolateral periaqueductal grey (VLPAG), lateral hypothalamus, paraventricular thalamus, caudate, insula, and area 24 in the cerebral cortex. CO_2_ exposure in rats increased number of c-fos**^+^** cells in the VLPAG (Johnson et al. 2011) and accordingly VLPAG neurons can a modulate the hypercapnic ventilatory response while having no effect on mean arterial pressure, heart rate, or body temperature (Lopes et al. 2012). Additionally,. In the lateral hypothalamus, the neuropeptide orexin is known to be involved in breathing behavior. Following mouse exposure to CO_2_ an increased c-fos expression in orexin neurons also reported (Sunanaga et al. 2009). In a study investigating c-fos expression in the paraventricular thalamus after myocardial infarction, a high level of c-fos**^+^** neurons were observed in this region. The authors interpreted this increase in c-fos to be associated with stress in the brain that may occur prior to irregular sympathetic response, such as heart rate, leading to myocardial infarction (Berquin et al. 2000). As part of the basal ganglia network, caudate nucleus has substantial connections with other basal ganglia nuclei such as the ventral tegmental area (VTA) (Brittain and Brown 2014). In mice, CO_2_ was found to modulate activity of GABAergic neurons within the VTA (Hill et al. 2020). Although we did not find CO_2_-induced increase of c-fos activity in marmoset VTA, we found CO_2_-activated cells in the closely connected caudate nucleus (Figure 1). These data suggest that changes in blood *P*CO_2_ could potentially regulate motor activity and/or reward behavior.

While previous research has detailed brainstem regions involved in chemosensation, much less is known of the involvement in cortical regions. We found two regions (namely, insula cortex and anterior cingulate cortex) that show CO_2_-dependent increases in c-fos expression in marmoset. Recently, it was suggested that insula might be involved in regulation of breathing behaviors (Golestani and Chen 2020) and that changes in blood CO_2_ levels are sensed not only by brainstem chemosensitive regions but also by some cortical regions, including insula (Parsons et al. 2001). Changes in area 24 activity, recorded via deep brain electrodes in humans, was observed following inhalation of varying percentages of CO_2_ (Holton et al. 2021). These data suggest that insula and area 24 cortical regions may be involved in interoception of breathing.

### 4.3 Phox2b and putative RTN region

Phox2b immunostaining has been used to identify RTN region in experimental animals (Ruffault et al. 2015; Kang et al. 2007; Abbott et al. 2011). Therefore, to identify CO_2_-activated Phox2b-expressing neurons (Kumar et al. 2015; Stornetta et al. 2006), we used immunostaining of both Phox2b and c-fos in marmosets exposed to 6% CO_2_ and mapped the Phox2b^+^ c-fos^+^ cells ventral to the facial nucleus, investigating the putative location of the RTN. Similar to other regions in the ventral brainstem, there was a basal level of c-fos activity within the RTN region in the control animals (animals not expose to acute hypercapnia) which could be related to the involvement of Phox2b^+^ neurons in modulation of breathing activity (Dereli et al. 2019; Fu et al. 2019; Fu et al. 2017; Tian et al. 2021). However, the number of Phox2b**^+^** c-fos**^+^** cells in parafacial region were increased by 10 folds in marmosets exposed to 6% CO_2_ (Figure 2). Our analysis of Phox2b**^+^** c-fos**^+^**cells indicated that there are ∼ 2500 neurons in the marmoset putative RTN region. This number is comparable to the number of RTN neurons in rats (∼ 2000 neurons) whereas the number of neurons in the macaque RTN is estimated as ∼2500 (Stornetta et al. 2006, Levy et al. 2019). This helps elucidate similarities in RTN size, density, and potential location as the rodent model is a similar physical size to the marmoset. This is comparable to a larger non-human primate such as a macaque where the tentative outline of the RTN matches reasonably well with the marmoset map of the RTN region based on Phox2b and c-fos expression patterns, however the number of Phox2B positive neurons in the RTN is much greater (Shi et al. 2017; Feldman and Del Negro 2006; Guyenet et al. 2010; Smith et al. 1989; Levy et al. 2019).

### 4.4 TPH and putative medullary raphé nuclei

Aside from the Phox2b**^+^** neuronal population in the RTN, the 5-HT neurons of the medullary 10aphe are also proposed to act as central CO_2_ chemosensor in rodents (Richerson et al. 2001; Richerson 2004; Nattie et al. 2004; Hodges and Richerson 2010; Corcoran et al. 2009; Ray et al. 2011; Bhandare et al. 2020). Similar to data in rodents, we found that a subset of serotonergic neurons in the brainstem are activated by systemic hypercapnia (Figure 3). Studies in rodents suggests that medullary 10aphe 5-HT neurons modulate ventilatory response to hypercapnia, however, DRN neurons are critical for arousal response to CO_2_ (Smith et al. 2018; Hodges and Richerson 2010). Using TPH and c-fos immunostaining, the putative location of the medullary 11aphe and dorsal 11aphe nucleus (DRN) was identified.

### 4.5 Brainstem Astrocytes

Recently data has implicated medullary astrocytes in central CO_2_ chemosensory responses (Turovsky et al. 2015; Huckstepp and Dale 2011; Beltrán-Castillo et al. 2017; Gourine et al. 2010; Sheikhbahaei, Turovsky, et al. 2018; Marina et al. 2018) (Turk et al. 2022; [Gourine and Dale 2022](https://sciwheel.com/work/signin?targetUrl=%2Fwork%2F%23%2Fitems)). Within the RTN region, astrocytes modulate breathing via pH-dependent release of ATP (Gourine et al. 2010). RTN astrocytes are morphologically different from other brainstem astrocytes; they have intermingled long processes that make a meshwork in the ventral surface of the medulla in rodents (Sheikhbahaei, Morris, et al. 2018). Interestingly, in rodents, a portion of RTN astrocytes express Phox2b (Czeisler et al. 2019). Consistent with these data, we observed long, and network-like processes of RTN astrocytes in the marmoset brain (Figure 4). In addition, our data suggest that the majority of astrocytic processes in marmoset’s RTN are Phox2b**^+^** (Figure 5). The location of these Phox2b**^+^** astrocytes (close to the ventral surface) together with the morphological similarities of marmoset astrocytes to rodent astrocytes in the RTN region, further strengthen the hypothesis that astrocytes in the ventral lateral medulla can act as central chemosensor in primates. However, more experiments are needed to directly define the chemomodulatory role of these astrocytes in control of breathing behaviors in primates.

As a non-human primate, the common marmoset has been proposed as a powerful animal model for social and physiological studies in neuroscience (Prins et al. 2017; Walker et al. 2017; Mitchell and Leopold 2015; Miller et al. 2016; Bishop et al. 2021; Mansfield 2003; Burkart and Finkenwirth 2015; SheikhBahaei 2020). Compared to rodents’ brain, the marmosets brain is more alike to that of humans (Passingham et al. 2013), however, the size of the marmoset - closer to that of a rodent – makes it an appealing laboratory animal model for physiological experiments. While the marmoset has the potential to be a compelling model to investigate a variety of neurophysiological functions, the brainstem nuclei of this animal have not been properly mapped. In this study, we mapped the central chemosensitive regions of RTN and medullary raphé in the brainstem. Interestingly, the location of RTN and raphé nuclei mapped in our study are similar to the location of these nuclei in rat, a similar size rodent. As the brainstem is a complex brain region with many unqiue functions and regions, more studies will be required to further map the other specialized respiratory nuclei in the marmoset brainstem.

## Conflict of interest

The authors declare no competing financial interests.

## Acknowledgements

We thank NINDS Light Microscopy Core (Director: Dr. Carolyn Smith), as well as NIMH Systems Neuroscience Imaging Resource (SNIR; Drs. Ted Usdin (Director) and Sarah Williams) for technical assistances. Anatomical MRI scanning was carried out in the NIMH, NINDS, and NEI Neurophysiology Imaging Facility Core (Director: Dr. David Leopold), with special thanks to Drs. David Yu and Frank Ye for technical assistance. We thank Dr. J-F. Brunet for providing Phox2b antibody. We thank Drs. David Leopold (NIMH), Yogita Chudasama (NIMH), and Jeffrey Smith (NINDS) for mentorship, support, and discussion. We thank Dr. Alexander Gourine for comments on the previous version of the manuscript. This work was supported by the Intramural Research Program of the NIH, NINDS and NIMH (ZIA NS009420 to SSB).

## Notes

### Competing Interest Statement

The authors have declared no competing interest.

